# Migration distance and mating system are not associated with genetic diversity and differentiation among bats (Chiroptera)

**DOI:** 10.1101/2023.09.28.559949

**Authors:** Matt J. Thorstensen, Alicia M. Korpach, Evelien de Greef, Levi Newediuk, Chloé Schmidt, Colin J. Garroway

**Author notes:** co-first author (Matt J. Thorstensen, Alicia M. Korpach).

## Abstract

Genetic variation is critical for evolutionary responses to environmental change. Links between genetic variation and behavioural or life history traits may reveal how varied strategies influence evolutionary trends in speciation and adaptation. Traits associated with movement typically correlate with population genetic structure and could help predict populations’ vulnerability to geographic processes such as habitat fragmentation and disease spread. With their wide diversity in behaviours and ecologies, bats provide a useful testing ground for hypotheses about population structure related to species-specific movement patterns. We used a global sample of microsatellite data (*n*=233 sites from 17 bat species) associated with published studies to examine potential links between genetic variation and migration and mating strategies. The genetic measures we tested were population-specific differentiation, gene diversity, and allelic richness. Using Bayesian models that accounted for phylogenetic distances among species, we identified no correlations between migration or mating strategy and genetic variation. Our results do not support long-standing hypotheses about dispersal-mediated genetic structure, and contrast with prior studies on bat genetic diversity and differentiation. We discuss the need for continued research into the complex association of ecological, biogeographical, and behavioural factors that facilitate gene flow among populations, especially in species with diverse movement patterns.

## Introduction

While genetic variation is required for evolutionary responses to environmental change and thus population resilience, its connections to behavioural and life history and traits are less clear (Lande, 1980; Mitton & Lewis, 1989; Hamrick & Godt, 1996; Barrett & Schluter, 2008; Schluter & Conte, 2009). These connections may underscore evolutionary processes in population and species-level divergence, as phenotypic variation may contribute to population isolation and ecological speciation (Lande, 1980; Rundle & Nosil, 2005). Behavioural traits related to motility are fundamental to most animals’ life histories, and how, when, and where they move during mating and dispersal directs gene flow patterns across the landscape (Cushman & Lewis, 2010) (Cushman and Lewis 2010). For example, geographical, ecological, and behavioural barriers to movement restrict gene flow and can generate structured populations in bats (Moussy *et al*., 2013). Dispersal ability (i.e., the ability to fly, swim, or move across potential barriers) is generally negatively correlated with genetic structure because reproductive individuals from populations of highly motile species are more able to interact and exchange genetic material (Bohonak, 1999; Bradbury *et al*., 2008; Medina *et al*., 2018). However, dispersal ability alone has inconsistent correlations with genetic structure among different species (Taylor *et al*., 2012; Burns & Broders, 2014). With widely-available genetic and ecological data, potential reciprocal links between movement-related traits and genetic variation are becoming more tractable for study across broadly distributed groups of organisms (Schmidt *et al*., 2020).

As the only flying mammals, bats are an interesting group to study connections between movement-related traits and genetic variation. Migration, colonization, and mating systems are often correlated with population structure in bats (Castella & Ruedi, 2000; Burland & Worthington Wilmer, 2001; Kerth & Morf, 2004; Furmankiewicz & Altringham, 2007). However, individual genetic studies are often difficult to compare because of varying marker types and measures of genetic variation (Moussy *et al*., 2013). A standardized examination of mating systems, migration strategy, and genetic variation can provide insight into how behavioural traits interact with genetic variation in bat species or populations. Bat species worldwide face threats related to climate change, habitat loss, overhunting, and novel fungal pathogens (Lorch *et al*., 2011; Frick, Kingston, & Flanders, 2020). Defining the traits that best predict genetic structure can help decision-makers identify groups of bats that may be at greater risk of loss of genetic variation, thereby requiring enhanced monitoring or protection.

One such movement trait that may be related to genetics is migration, the round-trip, seasonal movement of organisms between locations, and is a trait exhibited by numerous taxa to avoid seasonally harsh environments or exploit food resources (Shaw, 2016). It is widely accepted as adaptive, and long-distance migration patterns shaped by competition likely co-evolved with other life history traits (Alerstam, Hedenström, & Åkesson, 2003). Migratory movements have been related to increased genetic diversity in taxa as diverse as butterflies (Garcí ▫ Berro *et al*., 2023), mammals (Gustafson *et al*., 2017), and fish (Kovach, Gharrett, & Tallmon, 2013), making migration a potentially important factor in shaping population genetic structure.

A species’ mating system is another trait that depends on movement patterns (Shuster, 2009). Random or promiscuous mating reduces relatedness in family groups and increases genetic diversity (McCauley & O’Donnell, 1984; Gohli *et al*., 2013). In contrast, monogamy or polygamy theoretically decreases genetic variation in populations, reducing effective population size (Briton *et al*., 1994). However, work in wild bat populations showed that polygyny may increase genetic variation (Pérez-González, Mateos, & Carranza, 2009; Garg *et al*., 2012). Explanations for this inconsistency vary, but correlates of harem mating in some polygynous species (e.g., greater male-male competition) may promote increased genetic variation (Pérez-González *et al*., 2009; Garg *et al*., 2012).

We reasoned that migratory bats (male or female) that travel to swarming sites to breed would have a higher likelihood of interacting with individuals from different populations. Therefore, we classified bat species into three movement classes (long-distance, regional, and non-migrant) to test an association between migration distance and genetic differentiation. We hypothesized that the weakest population-specific genetic differentiation (Weir & Goudet, 2017) would occur in migratory bats, due to increased dispersal ability and greater likelihood of mixing between populations, leading to increased gene flow. We also classified bat species into two mating classes to test an association between mating strategy and genetic differentiation. Bat species can have polygamous (harem-forming), promiscuous (swarming), or (rarely) monogamous mating strategies (McCracken & Wilkinson, 2000). We hypothesized that genetic structure would be stronger in harem-forming species than in swarming species because bats with harems theoretically have fewer mates in a smaller geographic area compared to those that travel to swarming sites. We also hypothesized lower genetic variation (gene diversity and allelic richness) in bat species with shorter migration distances and that use harems. To provide a genetic context for these models of population genetics and life history traits, we assessed differences in the three genetic measures (genetic differentiation, gene diversity, and allelic richness) at a species level.

## Methods

### Genetic Measurement Collection Methods

The microsatellite data used in the present genetic analyses were compiled in previously published work (Schmidt *et al*., 2020). They were originally published in multiple research articles (Buchalski, Chaverri, & Vonhof, 2014; Burns, Frasier, & Broders, 2014; Witsenburg *et al*., 2015b; Boston *et al*., 2015b; Johnson *et al*., 2015; Moussy *et al*., 2015b; Razgour *et al*., 2015b; Baird *et al*., 2015; Vesterinen *et al*., 2016b; Günther *et al*., 2016b; Afonso *et al*., 2017b; Cleary, Waits, & Finegan, 2017b; Davy *et al*., 2017; Lausen *et al*., 2019) and several online repositories (Buchalski, Chaverri, & Vonhof, 2013; Boston *et al*., 2015a; Burns, Frasier, & Broders, 2015; Moussy *et al*., 2015a; Razgour *et al*., 2015a; Witsenburg *et al*., 2015a; Baerwald & Barclay, 2016; Günther *et al*., 2016a; Johnson *et al*., 2016; Vesterinen *et al*., 2016a; Afonso *et al*., 2017a; Cleary, Waits, & Finegan, 2017a; Santos & Meyer, 2017; Davy *et al*., 2018; Lausen *et al*., 2018). In total, we attained data from 17 bat species, representing 233 populations and 8,095 individuals (Table S1). We used all individuals for gene diversity and allelic richness metrics, and due to one population for some groups, we used a subset of samples to estimate genetic differentiation which included 12 species (228 populations with 7,618 individuals) (Figure 1; Table S2). Gene diversity was calculated with Adegenet v2.1.5 (Jombart, 2008), while genetic differentiation and allelic richness were calculated with hierfstat v0.5.7 (Goudet, 2005).

**Figure 1.**
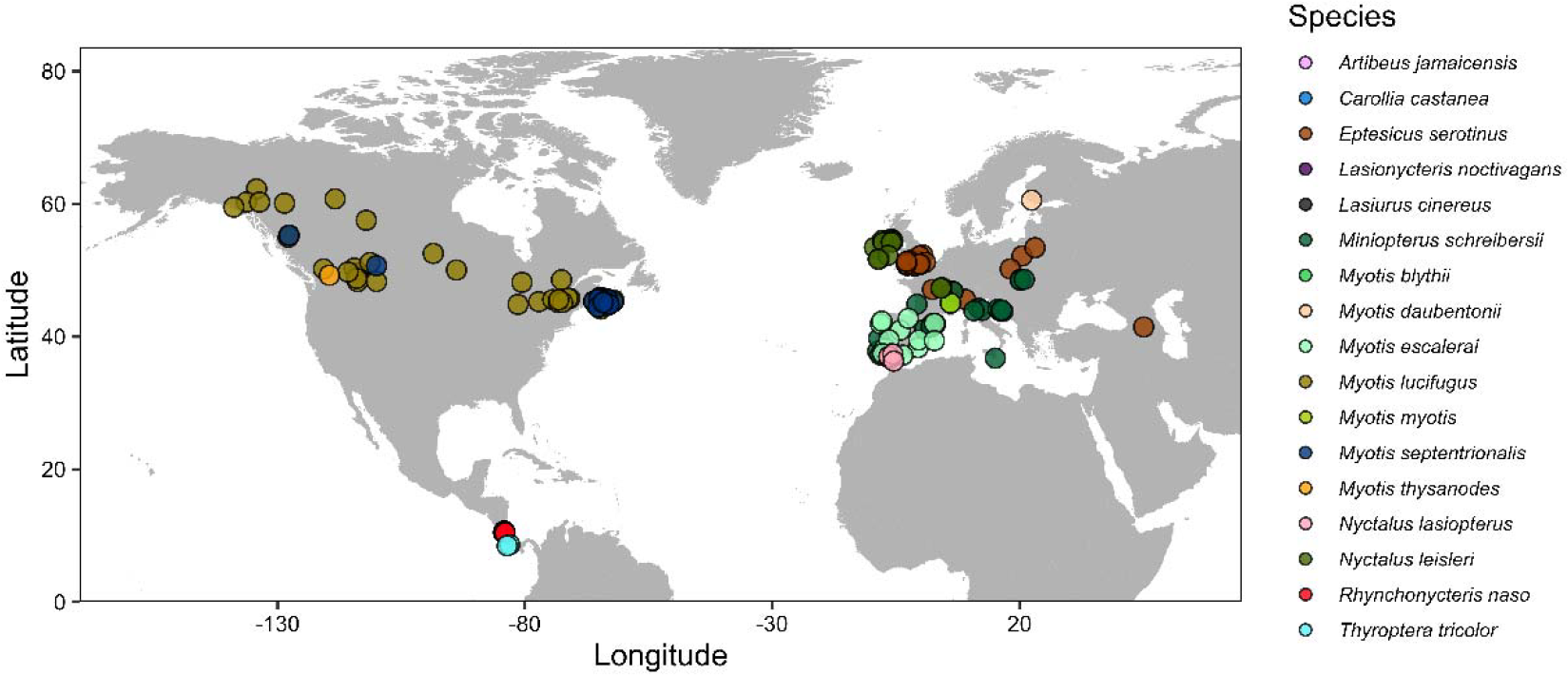
Map of bat genetic data and sampling locations used in the present analyses. Sampling locations are coloured by species. A total of 12 species, 228 sampling groups, and 7618 individuals were used for analyses of genetic differentiation (genetic differentiation) while 17 species, 233 sampling groups, and 8095 individuals were used for analyses of gene diversity (gene diversity) and allelic richness.

### Behaviour Classification Methods

Migration strategies were classified by species into non-migrants, regional migrants, and long-distance migrants based on their known life histories. Species observed to migrate between summer and winter roosts within the same region (movements may be tens to hundreds of km) were considered regional migrants, and migrants travelling between different regions (movements commonly greater than 1,000 km) were considered long-distance migrants. These categories roughly correspond to the distance categories used by Burns and Broders (2014). Similarly, where we could not find an explicit reference to a species’ migratory status, we classified them as non-migratory. Bat species were also classified by mating system, determined by whether or not they used harems in their reproduction. The full list of behaviour classifications is in Table S3, using several databases and research articles (Davis & Hitchcock, 1965; Fenton, 1969; Bradbury & Vehrencamp, 1977; Nagorsen & Brigham, 1993; Leu, 2000; Caceres & Barclay, 2000; Anderson, 2002; Vingiello, 2002; Wang, 2002; Keinath, 2004; Wohlgemuth *et al*., 2004; McGuire *et al*., 2012; Boston *et al*., 2012; Moussy *et al*., 2013; Cryan, Stricker, & Wunder, 2014; Ibáñez & Juste, 2016; Bentley, 2017; Fraser, Brooks, & Longstaffe, 2017; Taylor & Tuttle, 2019; Godlevska, Gazaryan, & Kruskop, 2021; Elliott, 2022; GBIF Secretariat, 2022a,b,c,d; Encyclopedia of Life, 2023; Jarso, 2023; NBN Atlas, 2023; UNEP/EUROBATS Secretariat, 2023).

### Population Genetics Across Species

Prior to modeling correlations between population genetic measures and behaviours, we examined differences in population genetic measures among species without modeling other variables. In the statistical computing environment R v4.2.1, the package brms v2.18.0 was used for Bayesian modeling, while tidyverse v2.0.0, tidybayes v3.0.3, ggbeeswarm v0.7.1, and patchwork v1.1.2 were useful for data management and model visualization (Bürkner, 2018; Wickham *et al*., 2019; Pedersen, 2022; R Core Team, 2022; Kay, 2023). We first examined differences in population genetic measures across species by fitting a series of models with species identity as a fixed effect.

The three population genetic measures used were genetic differentiation, gene diversity (also called expected heterozygosity), and allelic richness (Nei & Chesser, 1983; Weir & Goudet, 2017). Each Bayesian model was fit with a Gaussian distribution, using four independent Hamiltonian Monte Carlo chains across 5,000 warm-up and 15,000 sampling iterations each (60,000 sampling iterations total). For the models analyzing genetic differentiation or gene diversity, normal priors of mean 0 and standard deviation 1 were used for global intercepts and model coefficients. For the model analyzing allelic richness, we used normal priors of mean 0 and standard deviation 5. In all models, adapt delta was raised to 0.99 and maximum tree depth raised to 16. Model fit was assessed with the potential scale reduction statistic *R□*, visual inspection of trace plots, and visual inspection of posterior predictive checks over 100 draws in each model. Models were accepted only if *R*□ was 1.00 for all parameters. The R package emmeans v1.8.4-1 (Searle, Speed, & Milliken, 1980) was used to both gather posterior draws for visualization and for estimating marginal means to compare 95% highest posterior density (HPD) in a pairwise manner between species.

### Genetic-Life History Trait Correlations

We used a Bayesian approach to model potential relationships between migration strategy or mating system, and population genetic measures. The same statistical programs and packages were used for these models as in the models of population genetics across species. Bayesian models were fit with the following model structure in R:

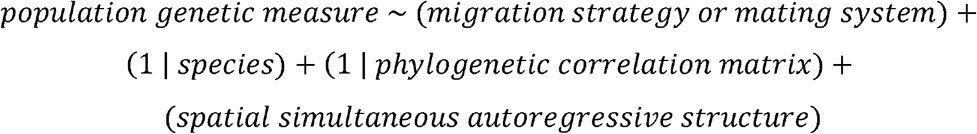

Here, genetic differentiation, gene diversity, and allelic richness were also used as population genetic measures. Thus, six models were run: three population genetic measures compared against either migration strategy or mating system.

Both migration strategy and mating systems were modeled as fixed effects. Species was included as a random intercept in each model to address factors independent of phylogenetic relatedness that may cause differences in species means for each genetic metric, such as niche or environmental effects (Hadfield & Nakagawa, 2010). A phylogenetic correlation matrix was also included as a random effect to address evolutionary relatedness among the bat species used in the present analyses. This phylogeny was created by using the R package taxize v0.9.00 to retrieve the taxonomic classifications and hierarchy for the list of species used against the National Center for Biotechnology Information database using the functions *classification* and *class2tree* (Figure S1) (Chamberlain & Szöcs, 2013). With this taxonomic tree, the R package ape v5.7 was used to calculate the phylogenetic correlation matrix used in each model with *vcv*.*phylo* (Paradis & Schliep, 2019). Model priors, distributions, sampling parameters, and contrasts were set and assessed with the same methods as in the models of population genetic measures across species.

To address the potential for spatial autocorrelation to affect model results, we used spatial autoregressive terms in brms with the *sar* function. Specifically, a K=4 nearest neighbors connection network of sampling locations was created with the R package adespatial v0.3-21. This network was used for specifying the spatial weighting matrix in the spatial simultaneous autoregressive structure, with type ‘lag’ chosen to model response variables.

We used the *hypothesis* function in brms to identify evidence ratios and posterior probabilities of differences between groups in several models with contrasts that were not significant in terms of HPD intervals. These models were run without global intercepts (unlike the overall models) to draw explicit comparisons among all variable levels, thus no prior was set for intercept, but priors remained the same for model coefficients (i.e., normal distributions with mean 0 and standard deviation 1 for genetic differentiation and gene diversity, and a normal distribution with mean 0 and standard deviation 5 for allelic richness). The models run in this manner were the genetic differentiation mating model to compare non-harem to harem mating strategies, and the gene diversity and allelic richness migration models, to quantify higher values for the long-distance migrants versus the non-migrants and regional migrants. Model fits were evaluated using the same metrics as the other models in the present study.

## Results

Across species, the serotine bat (*Eptesicus serotinus*) (Schreber, 1774) and common bent-wing bat (*Miniopterus schreibersii*) (Kuhl, 1817) tended to have higher genetic differentiation, and lower gene diversity and allelic richness than certain other species, such as the Jamaican fruit bat (*Artibeus jamaicensis*) (Leach, 1821) or chestnut short-tailed bat (*Carollia castanea*) (Allen, 1890) (Figure 2; Tables S4-S6). Overall, these results revealed species-specific differences in population-specific differentiation and genetic variation.

**Figure 2.**
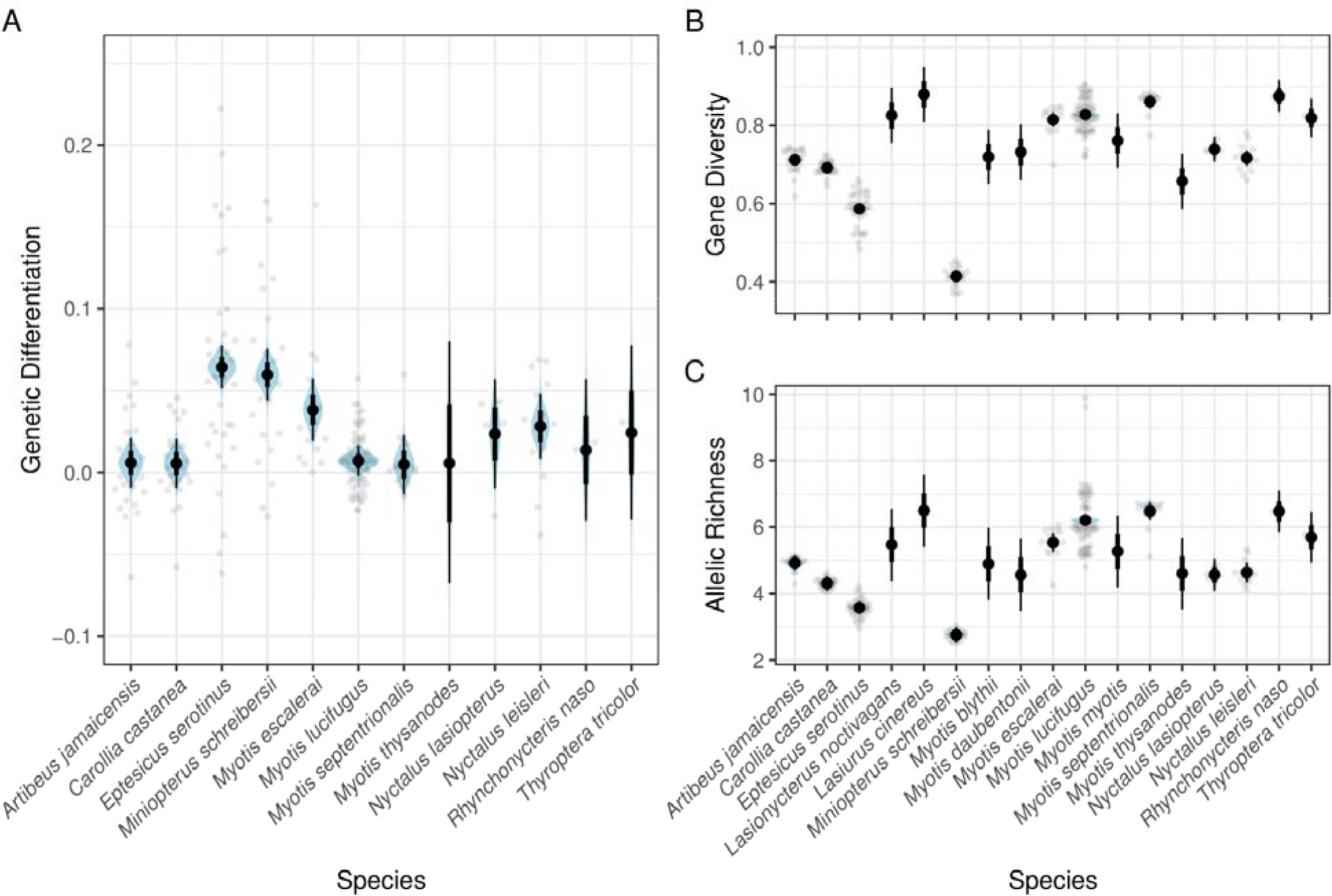
Estimates of population differentiation (genetic differentiation), gene diversity (gene diversity), and allelic richness across species. Posterior distributions are provided in blue, 95% credible intervals in thin black lines, and 66% credible intervals in bold black lines. Group-level data points used for modeling genetic measures across species are also provided.

We observed no differences in any comparison between migration or mating strategies using 95% HPD intervals (Figure 3; Tables S7-S8). Within the genetic differentiation model of mating strategies, the non-harem groups had a slightly higher genetic differentiation (0.01, -0.03 to 0.05 95% credible interval) than harem groups with an evidence ratio of 2.61 and posterior probability of 0.72 (Table S9). In the gene diversity model of migration strategies, long-distance migrants had higher gene diversity estimates than the regional and non-migrants with evidence ratios of 7.53and 1.86, and posterior probabilities of 0.88 and 0.65, respectively (Table S9). In the allelic richness model of migration strategies, long-distance migrants had higher allelic richness estimates than the regional and non-migrants with evidence ratios of 3.84 and 1.19, and posterior probabilities of 0.79 and 0.54, respectively (Table S9). We emphasize, however, that these differences were slight and none of the contrasts were ‘significant’ in the typical statistical sense of significance level α at 0.05.

**Figure 3.**
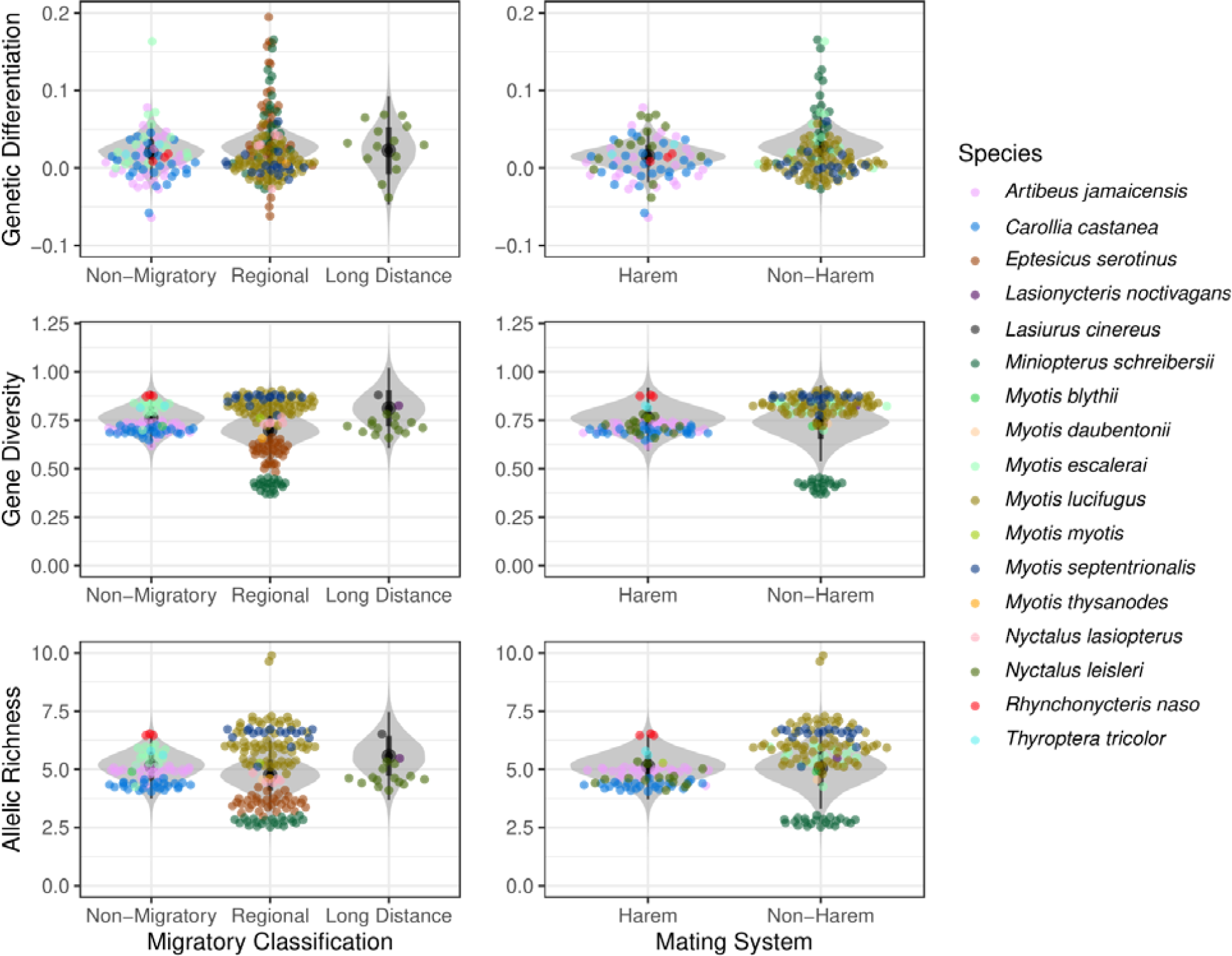
Estimates of population differentiation (genetic differentiation), gene diversity (gene diversity), and allelic richness compared between migration strategies (non-migratory, regional, long-distance) and mating systems (harm, non-harem) in bats. Posterior distributions are in gray, while group-level datapoints used for modeling genetic measures are coloured by species.

## Discussion

Despite species-level differences in population genetic measures, we did not find a relationship between migratory distance or mating system and population-specific differentiation and genetic variation among the bat populations studied. These results are consistent with work that identified high connectivity in bats over large regional scales, such as in the little brown bat (*Myotis lucifugus*) (Le Conte, 1831) (Burns *et al*., 2014). However, we had expected populations of species that exhibit larger dispersal distances to mix more, resulting in weaker population structure.

Our results did not support the hypothesis that long-distance migratory tendency results in genetic mixing. Wing morphology, which is correlated with migratory behaviour in birds, bats, and insects (Burns & Broders, 2014; Flockhart *et al*., 2017; Vincze *et al*., 2018), has emerged as a predictor of population structure in bats (Miller-Butterworth, Jacobs, & Harley, 2003; Olival, 2012; Burns & Broders, 2014). High wing loading allows fast flying with low maneuverability, and high wing aspect ratio allows efficient flights, which facilitates energy-efficient long-distance movements. However, long-distance migration dispersal (or the ability to do so) does not necessarily mean that homogeneous mixing of populations will occur. Regardless of the distance that a population migrates, a key assumption is that the migration movement results in breeding dispersal (Petit & Mayer, 2000); if strongly migratory species mate on the summer breeding grounds, as non-migrants do, their genetic structure would not be expected to differ from that of non-migrants’ (Moussy *et al*., 2013). It is possible that bat species with morphological features designed for use fast and efficient flights to mix far and wide on the breeding grounds, regardless of their migration behaviour. Studies that directly track migration movements, and discern the mating locations, of specific populations are required to reconcile the influences of morphological and behavioural traits.

Predicting genetic structure based on harem-forming and swarming behaviour may also be too simplistic, because a complex set of morphological, physiological, and behavioural traits combine with biogeographical features to impede or facilitate population mixing (Olival, 2012). Morphological characteristics such as brain size, testis size, and baculum length are correlated with female promiscuity or male fertilization success (Hosken & Stockley, 2004; Pitnick, Jones, & Wilkinson, 2006); all of which may contribute to shaping geographic genetic patterns among species. Some species of harem-forming bats exhibit high variation in levels of promiscuity (by both males and females) (Campbell, 2008; Garg *et al*., 2012), which could lead to weakened genetic structure among populations. Availability of food resources may influence short-term dispersal patterns on the breeding grounds; for instance frugivorous or nectivorous bats may need to move farther or more frequently than insectivorous bats, because insects are typically a more stable food source (Webb & Tidemann, 1996; Moreno-Valdez, Honeycutt, & Grant, 2004; Bontadina *et al*., 2008). Thus, for two species with similar mating strategies, frugivorous bats might incidentally have more genetic mixing because they are exposed to more individuals outside their populations while moving around to find food.

In addition to using relatively simplified models, some information on bats in the literature may not be accurate, leading to misclassification of the migratory or mating status for some of the species in our samples. Some bat species classified as non-migratory may move farther than we assume. Conversely, not all populations of species that we considered migratory actually migrate, which would result in a small average effect size in our analysis. For example, stable isotope analysis revealed high variation in the migratory movements of *Lasionycterus noctivagans* (Le Conte, 1831) (Fraser *et al*., 2017). Within species that are partially migratory, non-migratory populations can have greater genetic structure (e.g., *Eidolon helvum* (Kerr, 1792) (Juste, Ibáñez, & Machordom, 2000; Peel *et al*., 2013)). Even in species where all populations are migratory, those populations with low migratory connectivity can have greater genetic structure, possibly reflecting genetic isolation by distance (e.g., *M. lucifugus* (Vonhof, Russell, & Miller-Butterworth, 2015; Wilder, Kunz, & Sorenson, 2015)). While misclassification may have prevented us from detecting some genetic differences between migratory and non-migratory species, our null results also highlight the need for more studies on bat movement behaviour, as data on migration strategy is sparse among bats.

Several genetic reasons may underlie the present null results. Genetic variation was found to have a small to moderate influence on variance in migration timing in purple martins (*Progne subis*) (Linnaeus, 1758) (de Greef *et al*., 2023), and it is possible that migration and mating strategies are only weakly related to genetics in bats. Therefore, the population-level analyses conducted in the present study may have missed subtle, population-specific, and individual-level connections between the observed traits and genetics. Microsatellite data used for genetic differentiation is effective for testing overall patterns of genetic variation, it makes up small portions of the genome (Fischer *et al*., 2017). With increasing use of tools gaining larger representation of the genome (e.g., RAD-seq, whole genome sequencing), and improvements in methods to track bat movements, studying bat differentiation with increasing genome coverage may be useful for detecting linkages between movement-related life history traits and genetic variation. Alternatively, there may be no biological connection between the genetic variation and the observed traits in bats at the time these data were collected. Widespread phenotypic shifts may have induced prior shifts in gene flow or related processes in bats because of environmental change (Smeraldo *et al*., 2021), which may have altered the dynamics of how genotypes reflect phenotypes.

The differing sampling distribution and quantity across species in our study could affect the representation of different types of migration or mating systems. For example, some species were sampled more than others (ranging between 3104 bats across 66 sampling populations for *M. lucifugus*, and 6 bats across 1 sampling population for *M. thysanodes* (Miller, 1897)). This would change the spatial extent of the samples corresponding to each species’ respective ranges and artificially reduce genetic differentiation if few populations were sampled. We also assumed that distances moved during migration relative to range sizes did not affect the categorization of migratory strategies. Additionally, describing genetic structure in populations requires consideration of multiple seasons, such as summer maternity roosts, swarming sites, and hibernacula (Davy *et al*., 2015). Variation in the timing of sampling collection across species made it difficult to parse out seasonal effects on population structure overall, and we assumed samples came from discrete genetic populations. However, we can at least conclude that site-specific genetic differentiation and variation was not associated with migration or breeding strategy. These results show that bats potentially maintain genetic variation despite differences in behavioural strategies.

## Acknowledgements

We thank Quinn Fletcher for helpful conversations when designing this study. The present work would not have been possible without the work of numerous researchers in collecting, publishing, and making available the data used here. A Natural Sciences and Engineering Research Council of Canada Discovery Grant supported C.J.G.

## Data Availability Statement

All data used in the present study were downloaded from publicly available resources detailed in the methods, with specific repositories listed in the references.

