## Supplementary material for "Migration distance and mating system are not associated with genetic diversity and differentiation among bats (Chiroptera)": Table or Figure S

**Figure S1.** Phylogenetic tree of species used in the present study. Species names are coloured to be consistent with points used in other figures.

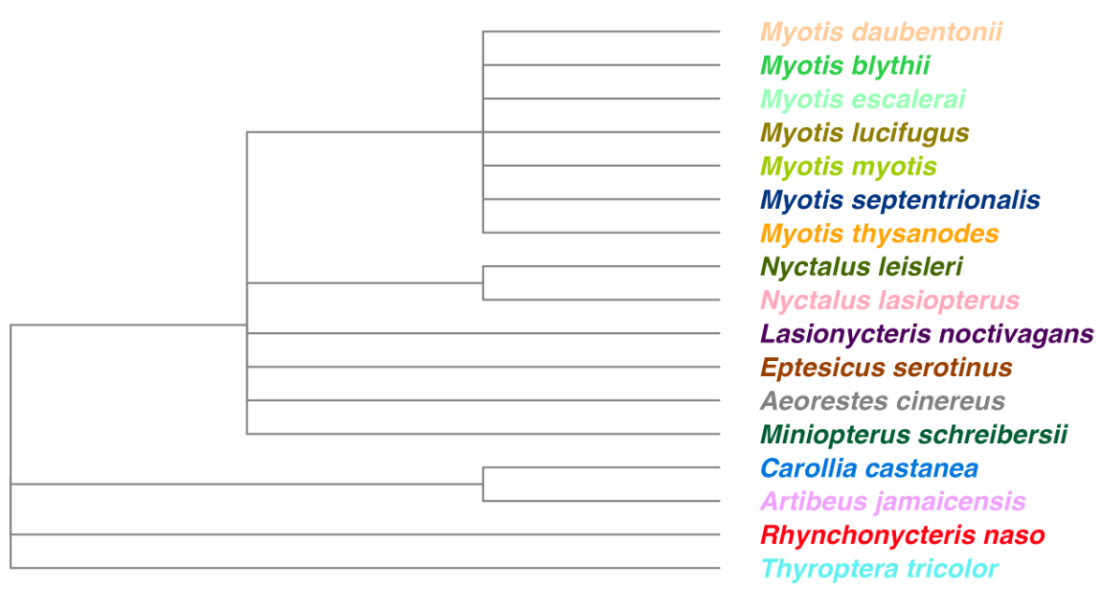

**Table S1**. Sample sizes of bat species by region used for analyses of gene diversity and allelic richness. Number of groups refers to the number of populations or groups that were present for a given species.

| **Region** | **Species** | **Number of Individuals** | **Number of Groups** |
| --- | --- | --- | --- |
| Asia | *Eptesicus serotinus* | 15 | 1 |
| Europe | *Eptesicus serotinus* | 679 | 33 |
| Europe | *Miniopterus schreibersii* | 312 | 22 |
| Europe | *Myotis blythii* | 12 | 1 |
| Europe | *Myotis daubentonii* | 106 | 1 |
| Europe | *Myotis escalerai* | 442 | 15 |
| Europe | *Myotis myotis* | 140 | 1 |
| Europe | *Nyctalus lasiopterus* | 214 | 5 |
| Europe | *Nyctalus leisleri* | 183 | 14 |
| North America | *Lasionycteris noctivagans* | 87 | 1 |
| North America | *Lasiurus cinereus* | 132 | 1 |
| North America | *Myotis lucifugus* | 3104 | 66 |
| North America | *Myotis septentrionalis* | 954 | 17 |
| North America | *Myotis thysanodes* | 6 | 1 |
| South America | *Artibeus jamaicensis* | 386 | 24 |
| South America | *Carollia castanea* | 359 | 25 |
| South America | *Rhynchonycteris naso* | 198 | 3 |
| South America | *Thyroptera tricolor* | 766 | 2 |
| **Total** |  | **8095** | **233** |

**Table S2**. Sample sizes of bat species by region used for analyses of genetic differentiation. Number of groups refers to the number of populations or groups that were present for a given species.

| **Region** | **Species** | **Number of Individuals** | **Number of Groups** |
| --- | --- | --- | --- |
| Asia | *Eptesicus serotinus* | 15 | 1 |
| Europe | *Eptesicus serotinus* | 679 | 33 |
| Europe | *Miniopterus schreibersii* | 312 | 22 |
| Europe | *Myotis escalerai* | 442 | 15 |
| Europe | *Nyctalus lasiopterus* | 214 | 5 |
| Europe | *Nyctalus leisleri* | 183 | 14 |
| North America | *Myotis lucifugus* | 3104 | 66 |
| North America | *Myotis septentrionalis* | 954 | 17 |
| North America | *Myotis thysanodes* | 6 | 1 |
| South America | *Artibeus jamaicensis* | 386 | 24 |
| South America | *Carollia castanea* | 359 | 25 |
| South America | *Rhynchonycteris naso* | 198 | 3 |
| South America | *Thyroptera tricolor* | 766 | 2 |
| **Total** |  | **7618** | **228** |

**Table S3.** Genetic sample regions and locations, migratory strategy and mating system classes and respective sources.

| **Species** | **Region** | **Location** | **Migratory strategy** | **Migration reference(s)** | **Mating system** | **Mating reference(s)** |
| --- | --- | --- | --- | --- | --- | --- |
| *Artibeus jamaicensis* | S. America | Costa Rica | non-migratory | Taylor & Tuttle 2019 | harem | Taylor & Tuttle 2019 |
| *Carollia castanea* | S. America | Costa Rica | non-migratory* | Taylor & Tuttle 2019 | harem | Taylor & Tuttle 2019 |
| *Eptesicus serotinus* | Asia | Georgia | regional | Godlevska et al. 2021 | swarm | Elliot 2022 |
| *Lasionycteris noctivagans* | N. America | Canada/USA | long-distance | McGuire et al. 2012; Fraser et al. 2017 | swarm | Bentley 2017 |
| *Lasiurus cinereus* | N. America | Canada/USA | long-distance | Cryan et al. 2014 | swarm | Anderson 2002 |
| *Miniopterus schreibersii* | Europe | Spain | regional | Leu 2000 | swarm | Leu 2000; Moussy et al. 2013 |
| *Myotis blythii* | Europe | France | non-migratory | GBIF Secretariat 2022a | swarm | GBIF Secretariat 2022a |
| *Myotis daubentonii* | Europe | Finland | regional | GBIF Secretariat 2022b | swarm | GBIF Secretariat 2022b |
| *Myotis escalerai* | Europe | Spain | non-migratory | GBIF Secretariat 2022c | swarm | Ibáñez & Juste 2016 |
| *Myotis lucifugus* | N. America | Canada/USA | regional | Davis & Hitchcock 1965; Fenton 1969 | swarm | Taylor & Tuttle 2019 |
| *Myotis myotis* | Europe | France | regional | UNEP/ EUROBATS | swarm | Wang 2002 |
| *Myotis septentrionalis* | N. America | Canada/USA | regional | Nagorsen & Brigham 1993 | swarm | Caceres & Barclay 2000 |
| *Myotis thysanodes* | N. America | Canada/USA | regional | Keinath 2004; Vingiello 2002 | swarm* | Vingiello 2002 |
| *Nyctalus lasiopterus* | Europe | Spain | regional* | GBIF Secretariat 2022d; Ibáñez et al. 2009 | swarm* | GBIF Secretariat 2022d; Ibáñez et al. 2009 |
| *Nyctalus leisleri* | Europe | N. Ireland | long-distance | NBN Atlas 2023; Wohlgemuth et al. 2004 | harem | Encyclopedia of Life 2023; Boston et al. 2012 |
| *Rhynchonycteris naso* | S. America | Costa Rica | non-migratory | Bradbury and Vehrencamp 1977 | harem | Taylor & Tuttle 2019 |
| *Thyroptera tricolor* | S. America | Costa Rica | non-migratory | Jarso 2023 | harem* | Jarso 2023 |

**Table S4**. Results for pairwise 95% highest posterior density (HPD) intervals of genetic differentiation between bat species.

| **Contrast** | **Estimate** | **Lower.HPD** | **Upper.HPD** |
| --- | --- | --- | --- |
| Artibeus_jamaicensis - Carollia_castanea | 4.15E-04 | -0.0209619 | 0.0213321 |
| Artibeus_jamaicensis - Eptesicus_serotinus | -0.058387 | -0.0783088 | -0.0385433 |
| Artibeus_jamaicensis - Miniopterus_schreibersii | -0.05377625 | -0.075998 | -0.0321608 |
| Artibeus_jamaicensis - Myotis_escalerai | -0.03233705 | -0.056976 | -0.00817999 |
| Artibeus_jamaicensis - Myotis_lucifugus | -0.00115406 | -0.0190543 | 0.0162717 |
| Artibeus_jamaicensis - Myotis_septentrionalis | 0.001008955 | -0.0231416 | 0.0242023 |
| Artibeus_jamaicensis - Myotis_thysanodes | 4.16E-04 | -0.0766275 | 0.075886 |
| Artibeus_jamaicensis - Nyctalus_lasiopterus | -0.01760845 | -0.0538398 | 0.0197863 |
| Artibeus_jamaicensis - Nyctalus_leisleri | -0.0222642 | -0.0478783 | 0.00217134 |
| Artibeus_jamaicensis - Rhynchonycteris_naso | -0.007690085 | -0.0534009 | 0.0376725 |
| Artibeus_jamaicensis - Thyroptera_tricolor | -0.0184711 | -0.0731625 | 0.0372801 |
| Carollia_castanea - Eptesicus_serotinus | -0.058827095 | -0.0785162 | -0.03913637 |
| Carollia_castanea - Miniopterus_schreibersii | -0.05424649 | -0.07639966 | -0.03275291 |
| Carollia_castanea - Myotis_escalerai | -0.03272875 | -0.0564921 | -0.0079048 |
| Carollia_castanea - Myotis_lucifugus | -0.001569425 | -0.019344038 | 0.015812206 |
| Carollia_castanea - Myotis_septentrionalis | 5.88E-04 | -0.02270176 | 0.02420972 |
| Carollia_castanea - Myotis_thysanodes | 2.19E-05 | -0.07656405 | 0.07592369 |
| Carollia_castanea - Nyctalus_lasiopterus | -0.018054115 | -0.0549784 | 0.01832177 |
| Carollia_castanea - Nyctalus_leisleri | -0.022708855 | -0.04729372 | 0.00282243 |
| Carollia_castanea - Rhynchonycteris_naso | -0.008098685 | -0.0541271 | 0.03685491 |
| Carollia_castanea - Thyroptera_tricolor | -0.018826094 | -0.07378493 | 0.0368095 |
| Eptesicus_serotinus - Miniopterus_schreibersii | 0.0046377 | -0.0157407 | 0.0251129 |
| Eptesicus_serotinus - Myotis_escalerai | 0.0261181 | 0.0024722 | 0.04874589 |
| Eptesicus_serotinus - Myotis_lucifugus | 0.057234 | 0.041568534 | 0.07338481 |
| Eptesicus_serotinus - Myotis_septentrionalis | 0.05948221 | 0.037187 | 0.081464 |
| Eptesicus_serotinus - Myotis_thysanodes | 0.058831165 | -0.0162777 | 0.1353074 |
| Eptesicus_serotinus - Nyctalus_lasiopterus | 0.04084985 | 0.0050179 | 0.0765796 |
| Eptesicus_serotinus - Nyctalus_leisleri | 0.036166365 | 0.0126608 | 0.0602673 |
| Eptesicus_serotinus - Rhynchonycteris_naso | 0.05080118 | 0.0054002 | 0.0954059 |
| Eptesicus_serotinus - Thyroptera_tricolor | 0.0401717 | -0.0143901 | 0.0950435 |
| Miniopterus_schreibersii - Myotis_escalerai | 0.02150025 | -0.0031155 | 0.04624391 |
| Miniopterus_schreibersii - Myotis_lucifugus | 0.052638115 | 0.034392 | 0.07102019 |
| Miniopterus_schreibersii - Myotis_septentrionalis | 0.0548845 | 0.0313815 | 0.0791957 |
| Miniopterus_schreibersii - Myotis_thysanodes | 0.05403605 | -0.0224866 | 0.1307412 |
| Miniopterus_schreibersii - Nyctalus_lasiopterus | 0.03611557 | -4.50E-04 | 0.0731315 |
| Miniopterus_schreibersii - Nyctalus_leisleri | 0.031553945 | 0.0058439 | 0.05665379 |
| Miniopterus_schreibersii - Rhynchonycteris_naso | 0.0460419 | -7.83E-04 | 0.0916019 |
| Miniopterus_schreibersii - Thyroptera_tricolor | 0.03532174 | -0.0207554 | 0.090359 |
| Myotis_escalerai - Myotis_lucifugus | 0.031100678 | 0.009787785 | 0.05191028 |
| Myotis_escalerai - Myotis_septentrionalis | 0.03334981 | 0.00722063 | 0.05946598 |
| Myotis_escalerai - Myotis_thysanodes | 0.032591225 | -0.0436679 | 0.1108173 |
| Myotis_escalerai - Nyctalus_lasiopterus | 0.0146844 | -0.0235297 | 0.05366522 |
| Myotis_escalerai - Nyctalus_leisleri | 0.01007275 | -0.0176487 | 0.03724574 |
| Myotis_escalerai - Rhynchonycteris_naso | 0.024614005 | -0.0226813 | 0.0711906 |
| Myotis_escalerai - Thyroptera_tricolor | 0.01397255 | -0.0421826 | 0.0709644 |
| Myotis_lucifugus - Myotis_septentrionalis | 0.0021519 | -0.01783442 | 0.022552356 |
| Myotis_lucifugus - Myotis_thysanodes | 0.00158221 | -0.07379796 | 0.07666731 |
| Myotis_lucifugus - Nyctalus_lasiopterus | -0.016412945 | -0.04998048 | 0.0194819 |
| Myotis_lucifugus - Nyctalus_leisleri | -0.0210647 | -0.04318452 | 8.19E-04 |
| Myotis_lucifugus - Rhynchonycteris_naso | -0.006567055 | -0.04937851 | 0.03887529 |
| Myotis_lucifugus - Thyroptera_tricolor | -0.017126445 | -0.07150046 | 0.03657629 |
| Myotis_septentrionalis - Myotis_thysanodes | -6.61E-04 | -0.076054247 | 0.07736722 |
| Myotis_septentrionalis - Nyctalus_lasiopterus | -0.018716665 | -0.056834937 | 0.01953197 |
| Myotis_septentrionalis - Nyctalus_leisleri | -0.023218113 | -0.0503073 | 0.00323552 |
| Myotis_septentrionalis - Rhynchonycteris_naso | -0.00879602 | -0.0557576 | 0.03794494 |
| Myotis_septentrionalis - Thyroptera_tricolor | -0.01935494 | -0.07402179 | 0.03863808 |
| Myotis_thysanodes - Nyctalus_lasiopterus | -0.018109795 | -0.1004088 | 0.06315047 |
| Myotis_thysanodes - Nyctalus_leisleri | -0.022762415 | -0.0983566 | 0.0561091 |
| Myotis_thysanodes - Rhynchonycteris_naso | -0.00828925 | -0.096558 | 0.076158 |
| Myotis_thysanodes - Thyroptera_tricolor | -0.018605075 | -0.1123054 | 0.0708778 |
| Nyctalus_lasiopterus - Nyctalus_leisleri | -0.0047254 | -0.042746483 | 0.03522353 |
| Nyctalus_lasiopterus - Rhynchonycteris_naso | 0.0097233 | -0.0447729 | 0.0639392 |
| Nyctalus_lasiopterus - Thyroptera_tricolor | -6.44E-04 | -0.0631854 | 0.0623163 |
| Nyctalus_leisleri - Rhynchonycteris_naso | 0.014539255 | -0.0327366 | 0.0624759 |
| Nyctalus_leisleri - Thyroptera_tricolor | 0.0038378 | -0.05192491 | 0.06171998 |
| Rhynchonycteris_naso - Thyroptera_tricolor | -0.010572285 | -0.0795568 | 0.0568948 |

**Table S5**. Results for pairwise 95% highest posterior density (HPD) intervals of gene diversity between bat species.

| **Contrast** | **Estimate** | **Lower.HPD** | **Upper.HPD** |
| --- | --- | --- | --- |
| Artibeus_jamaicensis - Carollia_castanea | 0.02036675 | 3.36E-04 | 0.0404187 |
| Artibeus_jamaicensis - Eptesicus_serotinus | 0.12524 | 0.106311 | 0.143834 |
| Artibeus_jamaicensis - Lasionycteris_noctivagans | -0.1136295 | -0.18509 | -0.041192 |
| Artibeus_jamaicensis - Lasiurus_cinereus | -0.16746 | -0.240031 | -0.0973007 |
| Artibeus_jamaicensis - Miniopterus_schreibersii | 0.2980095 | 0.276989 | 0.318414 |
| Artibeus_jamaicensis - Myotis_blythii | -0.00737293 | -0.0787351 | 0.0634933 |
| Artibeus_jamaicensis - Myotis_daubentonii | -0.0202908 | -0.0911911 | 0.053251 |
| Artibeus_jamaicensis - Myotis_escalerai | -0.102709 | -0.125491 | -0.0793511 |
| Artibeus_jamaicensis - Myotis_lucifugus | -0.115594 | -0.131968 | -0.0987542 |
| Artibeus_jamaicensis - Myotis_myotis | -0.04928735 | -0.120059 | 0.0220983 |
| Artibeus_jamaicensis - Myotis_septentrionalis | -0.149002 | -0.171468 | -0.126965 |
| Artibeus_jamaicensis - Myotis_thysanodes | 0.0549283 | -0.0163111 | 0.126715 |
| Artibeus_jamaicensis - Nyctalus_lasiopterus | -0.0275144 | -0.062339 | 0.00687103 |
| Artibeus_jamaicensis - Nyctalus_leisleri | -0.00465816 | -0.028934 | 0.0184455 |
| Artibeus_jamaicensis - Rhynchonycteris_naso | -0.162755 | -0.206286 | -0.119893 |
| Artibeus_jamaicensis - Thyroptera_tricolor | -0.1074015 | -0.158652 | -0.0555154 |
| Carollia_castanea - Eptesicus_serotinus | 0.1049699 | 0.0864486 | 0.12350292 |
| Carollia_castanea - Lasionycteris_noctivagans | -0.1338477 | -0.2054588 | -0.0617483 |
| Carollia_castanea - Lasiurus_cinereus | -0.187638 | -0.2585062 | -0.1154745 |
| Carollia_castanea - Miniopterus_schreibersii | 0.2776435 | 0.2566132 | 0.29781549 |
| Carollia_castanea - Myotis_blythii | -0.02772012 | -0.0983553 | 0.04335066 |
| Carollia_castanea - Myotis_daubentonii | -0.04041929 | -0.1124825 | 0.03162961 |
| Carollia_castanea - Myotis_escalerai | -0.1230655 | -0.1461205 | -0.1001886 |
| Carollia_castanea - Myotis_lucifugus | -0.1359423 | -0.1523438 | -0.1196033 |
| Carollia_castanea - Myotis_myotis | -0.0696154 | -0.1415457 | 9.19E-04 |
| Carollia_castanea - Myotis_septentrionalis | -0.1693119 | -0.1917858 | -0.147224 |
| Carollia_castanea - Myotis_thysanodes | 0.03459106 | -0.03552826 | 0.1074345 |
| Carollia_castanea - Nyctalus_lasiopterus | -0.04778375 | -0.0827965 | -0.01379324 |
| Carollia_castanea - Nyctalus_leisleri | -0.025032825 | -0.0482915 | -0.0012558 |
| Carollia_castanea - Rhynchonycteris_naso | -0.18314095 | -0.2263056 | -0.1396528 |
| Carollia_castanea - Thyroptera_tricolor | -0.12764675 | -0.1778785 | -0.0754974 |
| Eptesicus_serotinus - Lasionycteris_noctivagans | -0.2386845 | -0.309106 | -0.1662189 |
| Eptesicus_serotinus - Lasiurus_cinereus | -0.2927645 | -0.362926 | -0.2213372 |
| Eptesicus_serotinus - Miniopterus_schreibersii | 0.1726745 | 0.15301 | 0.191635 |
| Eptesicus_serotinus - Myotis_blythii | -0.132597945 | -0.2031381 | -0.0621696 |
| Eptesicus_serotinus - Myotis_daubentonii | -0.1454673 | -0.217272 | -0.0744394 |
| Eptesicus_serotinus - Myotis_escalerai | -0.228008 | -0.250073 | -0.2066806 |
| Eptesicus_serotinus - Myotis_lucifugus | -0.240897 | -0.2554 | -0.225808 |
| Eptesicus_serotinus - Myotis_myotis | -0.17465665 | -0.246859 | -0.10476932 |
| Eptesicus_serotinus - Myotis_septentrionalis | -0.2742615 | -0.29529 | -0.25358 |
| Eptesicus_serotinus - Myotis_thysanodes | -0.07020775 | -0.1406625 | 0.001328 |
| Eptesicus_serotinus - Nyctalus_lasiopterus | -0.15276773 | -0.1864415 | -0.1191573 |
| Eptesicus_serotinus - Nyctalus_leisleri | -0.129965769 | -0.1523651 | -0.10730807 |
| Eptesicus_serotinus - Rhynchonycteris_naso | -0.288161 | -0.329699 | -0.244875 |
| Eptesicus_serotinus - Thyroptera_tricolor | -0.2326075 | -0.28199 | -0.1804179 |
| Lasionycteris_noctivagans - Lasiurus_cinereus | -0.05379755 | -0.1514942 | 0.0472285 |
| Lasionycteris_noctivagans - Miniopterus_schreibersii | 0.4115335 | 0.3392744 | 0.483384 |
| Lasionycteris_noctivagans - Myotis_blythii | 0.10638665 | 0.0051949 | 0.2039061 |
| Lasionycteris_noctivagans - Myotis_daubentonii | 0.09319735 | -0.0045818 | 0.195232423 |
| Lasionycteris_noctivagans - Myotis_escalerai | 0.01084695 | -0.0632153 | 0.082767 |
| Lasionycteris_noctivagans - Myotis_lucifugus | -0.0021595 | -0.0733635 | 0.068629 |
| Lasionycteris_noctivagans - Myotis_myotis | 0.06411565 | -0.0347335 | 0.16313185 |
| Lasionycteris_noctivagans - Myotis_septentrionalis | -0.035499 | -0.1088657 | 0.036195 |
| Lasionycteris_noctivagans - Myotis_thysanodes | 0.168453685 | 0.06655882 | 0.265487 |
| Lasionycteris_noctivagans - Nyctalus_lasiopterus | 0.08613665 | 0.0079057 | 0.1615891 |
| Lasionycteris_noctivagans - Nyctalus_leisleri | 0.1087613 | 0.03563331 | 0.18136726 |
| Lasionycteris_noctivagans - Rhynchonycteris_naso | -0.0492935 | -0.1294886 | 0.033889 |
| Lasionycteris_noctivagans - Thyroptera_tricolor | 0.0061465 | -0.0803836 | 0.092028 |
| Lasiurus_cinereus - Miniopterus_schreibersii | 0.4653255 | 0.393565 | 0.536903 |
| Lasiurus_cinereus - Myotis_blythii | 0.1598726 | 0.0639207 | 0.2615484 |
| Lasiurus_cinereus - Myotis_daubentonii | 0.147203255 | 0.0492094 | 0.24927121 |
| Lasiurus_cinereus - Myotis_escalerai | 0.0645593 | -0.005588 | 0.13897 |
| Lasiurus_cinereus - Myotis_lucifugus | 0.0516655 | -0.0204946 | 0.120454 |
| Lasiurus_cinereus - Myotis_myotis | 0.118023 | 0.01975 | 0.2178649 |
| Lasiurus_cinereus - Myotis_septentrionalis | 0.018341 | -0.055573 | 0.088489 |
| Lasiurus_cinereus - Myotis_thysanodes | 0.22237095 | 0.1244254 | 0.321765 |
| Lasiurus_cinereus - Nyctalus_lasiopterus | 0.14002345 | 0.064247 | 0.2174734 |
| Lasiurus_cinereus - Nyctalus_leisleri | 0.1625269 | 0.0900461 | 0.2353042 |
| Lasiurus_cinereus - Rhynchonycteris_naso | 0.004488 | -0.076189 | 0.085654 |
| Lasiurus_cinereus - Thyroptera_tricolor | 0.059913 | -0.0263156 | 0.1459033 |
| Miniopterus_schreibersii - Myotis_blythii | -0.30534887 | -0.3770947 | -0.2352486 |
| Miniopterus_schreibersii - Myotis_daubentonii | -0.31809365 | -0.3903263 | -0.2459317 |
| Miniopterus_schreibersii - Myotis_escalerai | -0.4007326 | -0.423871 | -0.3768 |
| Miniopterus_schreibersii - Myotis_lucifugus | -0.413574 | -0.430854 | -0.3963562 |
| Miniopterus_schreibersii - Myotis_myotis | -0.3473505 | -0.4182466 | -0.2754863 |
| Miniopterus_schreibersii - Myotis_septentrionalis | -0.446903 | -0.46972 | -0.424287 |
| Miniopterus_schreibersii - Myotis_thysanodes | -0.2429387 | -0.3140622 | -0.171239 |
| Miniopterus_schreibersii - Nyctalus_lasiopterus | -0.32547175 | -0.360386 | -0.29016294 |
| Miniopterus_schreibersii - Nyctalus_leisleri | -0.302726445 | -0.32654891 | -0.27837894 |
| Miniopterus_schreibersii - Rhynchonycteris_naso | -0.4608275 | -0.503437 | -0.417056 |
| Miniopterus_schreibersii - Thyroptera_tricolor | -0.405341 | -0.456541 | -0.3533907 |
| Myotis_blythii - Myotis_daubentonii | -0.0126671 | -0.1126628 | 0.0846894 |
| Myotis_blythii - Myotis_escalerai | -0.095533647 | -0.168542 | -0.024694 |
| Myotis_blythii - Myotis_lucifugus | -0.108213435 | -0.1796042 | -0.0397107 |
| Myotis_blythii - Myotis_myotis | -0.042215056 | -0.14028528 | 0.057029 |
| Myotis_blythii - Myotis_septentrionalis | -0.14160471 | -0.2125416 | -0.0695417 |
| Myotis_blythii - Myotis_thysanodes | 0.06234425 | -0.0372112 | 0.1615647 |
| Myotis_blythii - Nyctalus_lasiopterus | -0.02005554 | -0.0971163 | 0.05422831 |
| Myotis_blythii - Nyctalus_leisleri | 0.0025036 | -0.0701405 | 0.0736618 |
| Myotis_blythii - Rhynchonycteris_naso | -0.155404105 | -0.2363139 | -0.0755345 |
| Myotis_blythii - Thyroptera_tricolor | -0.09978545 | -0.1876124 | -0.0173224 |
| Myotis_daubentonii - Myotis_escalerai | -0.0826772 | -0.1542245 | -0.0083168 |
| Myotis_daubentonii - Myotis_lucifugus | -0.0955065 | -0.1663966 | -0.024288 |
| Myotis_daubentonii - Myotis_myotis | -0.02956325 | -0.127856 | 0.0700561 |
| Myotis_daubentonii - Myotis_septentrionalis | -0.12894865 | -0.1996909 | -0.055043 |
| Myotis_daubentonii - Myotis_thysanodes | 0.075267535 | -0.0253312 | 0.1738028 |
| Myotis_daubentonii - Nyctalus_lasiopterus | -0.00724002 | -0.0834203 | 0.07140595 |
| Myotis_daubentonii - Nyctalus_leisleri | 0.01541192 | -0.0583603 | 0.0871538 |
| Myotis_daubentonii - Rhynchonycteris_naso | -0.14282225 | -0.2226059 | -0.059196 |
| Myotis_daubentonii - Thyroptera_tricolor | -0.08689685 | -0.1722405 | 2.93E-04 |
| Myotis_escalerai - Myotis_lucifugus | -0.01285785 | -0.0330899 | 0.007269 |
| Myotis_escalerai - Myotis_myotis | 0.05337735 | -0.0178965 | 0.1266347 |
| Myotis_escalerai - Myotis_septentrionalis | -0.04624695 | -0.0714241 | -0.021473 |
| Myotis_escalerai - Myotis_thysanodes | 0.1577597 | 0.084733 | 0.229265 |
| Myotis_escalerai - Nyctalus_lasiopterus | 0.07523815 | 0.0387602 | 0.11199125 |
| Myotis_escalerai - Nyctalus_leisleri | 0.09805 | 0.07118163 | 0.12346751 |
| Myotis_escalerai - Rhynchonycteris_naso | -0.0600791 | -0.105038 | -0.015135 |
| Myotis_escalerai - Thyroptera_tricolor | -0.00457015 | -0.059152 | 0.0468324 |
| Myotis_lucifugus - Myotis_myotis | 0.06633455 | -0.003427 | 0.1375098 |
| Myotis_lucifugus - Myotis_septentrionalis | -0.0333505 | -0.052491 | -0.014198 |
| Myotis_lucifugus - Myotis_thysanodes | 0.17065425 | 0.0997134 | 0.241562 |
| Myotis_lucifugus - Nyctalus_lasiopterus | 0.0881493 | 0.0549916 | 0.11997591 |
| Myotis_lucifugus - Nyctalus_leisleri | 0.110903325 | 0.0901367 | 0.1316021 |
| Myotis_lucifugus - Rhynchonycteris_naso | -0.0471985 | -0.088985 | -0.005568 |
| Myotis_lucifugus - Thyroptera_tricolor | 0.0082768 | -0.040832 | 0.0590309 |
| Myotis_myotis - Myotis_septentrionalis | -0.09956295 | -0.1704385 | -0.026516 |
| Myotis_myotis - Myotis_thysanodes | 0.104412855 | 0.0042328 | 0.2023958 |
| Myotis_myotis - Nyctalus_lasiopterus | 0.02188241 | -0.054955023 | 0.09787704 |
| Myotis_myotis - Nyctalus_leisleri | 0.044485355 | -0.0284272 | 0.11628283 |
| Myotis_myotis - Rhynchonycteris_naso | -0.11355305 | -0.1943097 | -0.0329054 |
| Myotis_myotis - Thyroptera_tricolor | -0.0581811 | -0.1432017 | 0.0269195 |
| Myotis_septentrionalis - Myotis_thysanodes | 0.2042171 | 0.1330388 | 0.276658 |
| Myotis_septentrionalis - Nyctalus_lasiopterus | 0.1215637 | 0.0860699 | 0.15801474 |
| Myotis_septentrionalis - Nyctalus_leisleri | 0.14427115 | 0.11933324 | 0.1702305 |
| Myotis_septentrionalis - Rhynchonycteris_naso | -0.0137395 | -0.057936 | 0.030262 |
| Myotis_septentrionalis - Thyroptera_tricolor | 0.041671 | -0.011252 | 0.0932325 |
| Myotis_thysanodes - Nyctalus_lasiopterus | -0.08230425 | -0.1605384 | -0.007089159 |
| Myotis_thysanodes - Nyctalus_leisleri | -0.05980087 | -0.1295108 | 0.01589597 |
| Myotis_thysanodes - Rhynchonycteris_naso | -0.2177058 | -0.299879 | -0.1376263 |
| Myotis_thysanodes - Thyroptera_tricolor | -0.16236655 | -0.247225 | -0.0766981 |
| Nyctalus_lasiopterus - Nyctalus_leisleri | 0.022780482 | -0.01387203 | 0.0594693 |
| Nyctalus_lasiopterus - Rhynchonycteris_naso | -0.1353482 | -0.1854981 | -0.0825934 |
| Nyctalus_lasiopterus - Thyroptera_tricolor | -0.079804 | -0.14122365 | -0.0229865 |
| Nyctalus_leisleri - Rhynchonycteris_naso | -0.15808835 | -0.2030501 | -0.1134028 |
| Nyctalus_leisleri - Thyroptera_tricolor | -0.10265657 | -0.15603027 | -0.05010746 |
| Rhynchonycteris_naso - Thyroptera_tricolor | 0.0555005 | -0.008732 | 0.1194362 |

**Table S6**. Results for pairwise 95% highest posterior density (HPD) intervals of allelic richness between bat species.

| **Contrast** | **Estimate** | **Lower.HPD** | **Upper.HPD** |
| --- | --- | --- | --- |
| Artibeus_jamaicensis - Carollia_castanea | 0.6155485 | 0.301002 | 0.920883 |
| Artibeus_jamaicensis - Eptesicus_serotinus | 1.34767 | 1.0665 | 1.64102 |
| Artibeus_jamaicensis - Lasionycteris_noctivagans | -0.5426325 | -1.63259 | 0.581457 |
| Artibeus_jamaicensis - Lasiurus_cinereus | -1.57705 | -2.67848 | -0.468876 |
| Artibeus_jamaicensis - Miniopterus_schreibersii | 2.16393 | 1.84177 | 2.48036 |
| Artibeus_jamaicensis - Myotis_blythii | 0.0269005 | -1.07579 | 1.13698 |
| Artibeus_jamaicensis - Myotis_daubentonii | 0.363178 | -0.754359 | 1.46806 |
| Artibeus_jamaicensis - Myotis_escalerai | -0.6179775 | -0.976037 | -0.259149 |
| Artibeus_jamaicensis - Myotis_lucifugus | -1.28962 | -1.54558 | -1.03205 |
| Artibeus_jamaicensis - Myotis_myotis | -0.3425905 | -1.45994 | 0.736838 |
| Artibeus_jamaicensis - Myotis_septentrionalis | -1.557445 | -1.89795 | -1.20929 |
| Artibeus_jamaicensis - Myotis_thysanodes | 0.3180535 | -0.790326 | 1.41206 |
| Artibeus_jamaicensis - Nyctalus_lasiopterus | 0.357481 | -0.179974 | 0.887884 |
| Artibeus_jamaicensis - Nyctalus_leisleri | 0.2879805 | -0.0860494 | 0.650552 |
| Artibeus_jamaicensis - Rhynchonycteris_naso | -1.553745 | -2.22359 | -0.890163 |
| Artibeus_jamaicensis - Thyroptera_tricolor | -0.767736 | -1.55633 | 0.0246097 |
| Carollia_castanea - Eptesicus_serotinus | 0.733287 | 0.4469 | 1.016319 |
| Carollia_castanea - Lasionycteris_noctivagans | -1.158783 | -2.277321 | -0.066096 |
| Carollia_castanea - Lasiurus_cinereus | -2.191834 | -3.280633 | -1.066651 |
| Carollia_castanea - Miniopterus_schreibersii | 1.5484205 | 1.237871 | 1.871796 |
| Carollia_castanea - Myotis_blythii | -0.58854565 | -1.6965 | 0.522629 |
| Carollia_castanea - Myotis_daubentonii | -0.253008 | -1.355442 | 0.863354 |
| Carollia_castanea - Myotis_escalerai | -1.2323895 | -1.591242 | -0.883941 |
| Carollia_castanea - Myotis_lucifugus | -1.9046455 | -2.165698 | -1.657848 |
| Carollia_castanea - Myotis_myotis | -0.95802 | -2.033102 | 0.171634 |
| Carollia_castanea - Myotis_septentrionalis | -2.173765 | -2.51562 | -1.836201 |
| Carollia_castanea - Myotis_thysanodes | -0.299091 | -1.407796 | 0.788992 |
| Carollia_castanea - Nyctalus_lasiopterus | -0.2593455 | -0.790153 | 0.271919 |
| Carollia_castanea - Nyctalus_leisleri | -0.326552 | -0.686794 | 0.041772 |
| Carollia_castanea - Rhynchonycteris_naso | -2.170154 | -2.822538 | -1.501303 |
| Carollia_castanea - Thyroptera_tricolor | -1.381542 | -2.175203 | -0.591007 |
| Eptesicus_serotinus - Lasionycteris_noctivagans | -1.8928105 | -2.98672 | -0.788133 |
| Eptesicus_serotinus - Lasiurus_cinereus | -2.9239 | -3.99621 | -1.798793 |
| Eptesicus_serotinus - Miniopterus_schreibersii | 0.81616 | 0.52011 | 1.10834 |
| Eptesicus_serotinus - Myotis_blythii | -1.32183525 | -2.438772 | -0.23101 |
| Eptesicus_serotinus - Myotis_daubentonii | -0.983949 | -2.09206 | 0.115 |
| Eptesicus_serotinus - Myotis_escalerai | -1.9646265 | -2.30181 | -1.628782 |
| Eptesicus_serotinus - Myotis_lucifugus | -2.636955 | -2.86461 | -2.40973 |
| Eptesicus_serotinus - Myotis_myotis | -1.69365 | -2.80547 | -0.626461 |
| Eptesicus_serotinus - Myotis_septentrionalis | -2.90555 | -3.22779 | -2.58488 |
| Eptesicus_serotinus - Myotis_thysanodes | -1.030247 | -2.10453 | 0.07914 |
| Eptesicus_serotinus - Nyctalus_lasiopterus | -0.991092 | -1.5139228 | -0.472104 |
| Eptesicus_serotinus - Nyctalus_leisleri | -1.059242 | -1.399676 | -0.713471 |
| Eptesicus_serotinus - Rhynchonycteris_naso | -2.901945 | -3.55304 | -2.25042 |
| Eptesicus_serotinus - Thyroptera_tricolor | -2.115271 | -2.88946 | -1.333185 |
| Lasionycteris_noctivagans - Lasiurus_cinereus | -1.0298065 | -2.5454532 | 0.5109934 |
| Lasionycteris_noctivagans - Miniopterus_schreibersii | 2.7109195 | 1.613544 | 3.82376 |
| Lasionycteris_noctivagans - Myotis_blythii | 0.57171 | -0.959336 | 2.134564 |
| Lasionycteris_noctivagans - Myotis_daubentonii | 0.911341 | -0.6325721 | 2.436187 |
| Lasionycteris_noctivagans - Myotis_escalerai | -0.0738565 | -1.170929 | 1.0708499 |
| Lasionycteris_noctivagans - Myotis_lucifugus | -0.7448445 | -1.836395 | 0.34984 |
| Lasionycteris_noctivagans - Myotis_myotis | 0.204618 | -1.354668 | 1.703982 |
| Lasionycteris_noctivagans - Myotis_septentrionalis | -1.013635 | -2.125809 | 0.10547 |
| Lasionycteris_noctivagans - Myotis_thysanodes | 0.86541405 | -0.6913716 | 2.36746 |
| Lasionycteris_noctivagans - Nyctalus_lasiopterus | 0.89951825 | -0.2972811 | 2.064983 |
| Lasionycteris_noctivagans - Nyctalus_leisleri | 0.8353425 | -0.320366 | 1.924924 |
| Lasionycteris_noctivagans - Rhynchonycteris_naso | -1.0092955 | -2.253405 | 0.243736 |
| Lasionycteris_noctivagans - Thyroptera_tricolor | -0.2238275 | -1.5697411 | 1.084143 |
| Lasiurus_cinereus - Miniopterus_schreibersii | 3.737595 | 2.621821 | 4.84342 |
| Lasiurus_cinereus - Myotis_blythii | 1.600094 | 0.07359 | 3.154033 |
| Lasiurus_cinereus - Myotis_daubentonii | 1.9376234 | 0.403807 | 3.481677 |
| Lasiurus_cinereus - Myotis_escalerai | 0.958379 | -0.169366 | 2.062189 |
| Lasiurus_cinereus - Myotis_lucifugus | 0.28527 | -0.80485 | 1.384884 |
| Lasiurus_cinereus - Myotis_myotis | 1.2296805 | -0.302799 | 2.751048 |
| Lasiurus_cinereus - Myotis_septentrionalis | 0.018415 | -1.108556 | 1.12456 |
| Lasiurus_cinereus - Myotis_thysanodes | 1.8946335 | 0.363112 | 3.41115 |
| Lasiurus_cinereus - Nyctalus_lasiopterus | 1.9291255 | 0.752777 | 3.139878 |
| Lasiurus_cinereus - Nyctalus_leisleri | 1.865 | 0.747655 | 2.994831 |
| Lasiurus_cinereus - Rhynchonycteris_naso | 0.01883 | -1.227357 | 1.27603 |
| Lasiurus_cinereus - Thyroptera_tricolor | 0.80529135 | -0.504877 | 2.14313739 |
| Miniopterus_schreibersii - Myotis_blythii | -2.1372105 | -3.218972 | -1.00857 |
| Miniopterus_schreibersii - Myotis_daubentonii | -1.80192 | -2.900739 | -0.67363 |
| Miniopterus_schreibersii - Myotis_escalerai | -2.781516 | -3.150225 | -2.429669 |
| Miniopterus_schreibersii - Myotis_lucifugus | -3.45374 | -3.71615 | -3.18435 |
| Miniopterus_schreibersii - Myotis_myotis | -2.5070365 | -3.61051 | -1.412178 |
| Miniopterus_schreibersii - Myotis_septentrionalis | -3.72208 | -4.07621 | -3.37404 |
| Miniopterus_schreibersii - Myotis_thysanodes | -1.845275 | -2.961317 | -0.75334 |
| Miniopterus_schreibersii - Nyctalus_lasiopterus | -1.807921 | -2.3430418 | -1.267885 |
| Miniopterus_schreibersii - Nyctalus_leisleri | -1.8752863 | -2.2475149 | -1.508174 |
| Miniopterus_schreibersii - Rhynchonycteris_naso | -3.71916 | -4.40683 | -3.06626 |
| Miniopterus_schreibersii - Thyroptera_tricolor | -2.9313905 | -3.72698 | -2.136804 |
| Myotis_blythii - Myotis_daubentonii | 0.337211 | -1.200685 | 1.874288 |
| Myotis_blythii - Myotis_escalerai | -0.64378975 | -1.754406 | 0.482484 |
| Myotis_blythii - Myotis_lucifugus | -1.3154677 | -2.42188 | -0.233671 |
| Myotis_blythii - Myotis_myotis | -0.37031565 | -1.907846 | 1.140122 |
| Myotis_blythii - Myotis_septentrionalis | -1.5852293 | -2.712244 | -0.485621 |
| Myotis_blythii - Myotis_thysanodes | 0.290418575 | -1.256451 | 1.802594 |
| Myotis_blythii - Nyctalus_lasiopterus | 0.330497 | -0.8482782 | 1.518109 |
| Myotis_blythii - Nyctalus_leisleri | 0.263404 | -0.889054 | 1.360467 |
| Myotis_blythii - Rhynchonycteris_naso | -1.5808665 | -2.84577 | -0.331013 |
| Myotis_blythii - Thyroptera_tricolor | -0.79454712 | -2.13167 | 0.5255302 |
| Myotis_daubentonii - Myotis_escalerai | -0.9821855 | -2.113273 | 0.133364 |
| Myotis_daubentonii - Myotis_lucifugus | -1.654757 | -2.73169 | -0.535275 |
| Myotis_daubentonii - Myotis_myotis | -0.707095 | -2.22813 | 0.832246 |
| Myotis_daubentonii - Myotis_septentrionalis | -1.9227925 | -3.03833 | -0.798614 |
| Myotis_daubentonii - Myotis_thysanodes | -0.048195 | -1.577069 | 1.4741818 |
| Myotis_daubentonii - Nyctalus_lasiopterus | -0.010432 | -1.196602 | 1.185134 |
| Myotis_daubentonii - Nyctalus_leisleri | -0.07584 | -1.195307 | 1.056213 |
| Myotis_daubentonii - Rhynchonycteris_naso | -1.9189245 | -3.19541 | -0.679684 |
| Myotis_daubentonii - Thyroptera_tricolor | -1.1314365 | -2.454195 | 0.19731 |
| Myotis_escalerai - Myotis_lucifugus | -0.673105 | -0.985118 | -0.363773 |
| Myotis_escalerai - Myotis_myotis | 0.2727195 | -0.840105 | 1.384136 |
| Myotis_escalerai - Myotis_septentrionalis | -0.9411035 | -1.318471 | -0.547697 |
| Myotis_escalerai - Myotis_thysanodes | 0.936931 | -0.178964 | 2.045975 |
| Myotis_escalerai - Nyctalus_lasiopterus | 0.9737334 | 0.421209 | 1.538805 |
| Myotis_escalerai - Nyctalus_leisleri | 0.9052055 | 0.498346 | 1.310431 |
| Myotis_escalerai - Rhynchonycteris_naso | -0.938751 | -1.620354 | -0.252107 |
| Myotis_escalerai - Thyroptera_tricolor | -0.1489818 | -0.955509 | 0.665449 |
| Myotis_lucifugus - Myotis_myotis | 0.945422 | -0.17635 | 1.992013 |
| Myotis_lucifugus - Myotis_septentrionalis | -0.26799 | -0.56024 | 0.02926 |
| Myotis_lucifugus - Myotis_thysanodes | 1.607222 | 0.501862 | 2.66762 |
| Myotis_lucifugus - Nyctalus_lasiopterus | 1.645642 | 1.1502954 | 2.163629 |
| Myotis_lucifugus - Nyctalus_leisleri | 1.577819 | 1.257911 | 1.899689 |
| Myotis_lucifugus - Rhynchonycteris_naso | -0.265725 | -0.9032 | 0.379876 |
| Myotis_lucifugus - Thyroptera_tricolor | 0.523413 | -0.24407 | 1.2994133 |
| Myotis_myotis - Myotis_septentrionalis | -1.2147225 | -2.307625 | -0.07872 |
| Myotis_myotis - Myotis_thysanodes | 0.66298765 | -0.869841 | 2.167171 |
| Myotis_myotis - Nyctalus_lasiopterus | 0.69867005 | -0.472919 | 1.887804 |
| Myotis_myotis - Nyctalus_leisleri | 0.630473 | -0.479679 | 1.743416 |
| Myotis_myotis - Rhynchonycteris_naso | -1.212554 | -2.479193 | 0.01077 |
| Myotis_myotis - Thyroptera_tricolor | -0.4270345 | -1.7551975 | 0.886071 |
| Myotis_septentrionalis - Myotis_thysanodes | 1.87582544 | 0.762789 | 2.96442 |
| Myotis_septentrionalis - Nyctalus_lasiopterus | 1.915108 | 1.3648603 | 2.47625 |
| Myotis_septentrionalis - Nyctalus_leisleri | 1.8461907 | 1.4458021 | 2.232966 |
| Myotis_septentrionalis - Rhynchonycteris_naso | 0.00371 | -0.68454 | 0.674369 |
| Myotis_septentrionalis - Thyroptera_tricolor | 0.7918115 | -0.01762 | 1.5834833 |
| Myotis_thysanodes - Nyctalus_lasiopterus | 0.0376075 | -1.1271888 | 1.2295133 |
| Myotis_thysanodes - Nyctalus_leisleri | -0.029221 | -1.147649 | 1.073729 |
| Myotis_thysanodes - Rhynchonycteris_naso | -1.871438 | -3.13335 | -0.636105 |
| Myotis_thysanodes - Thyroptera_tricolor | -1.0811845 | -2.40557 | 0.2081692 |
| Nyctalus_lasiopterus - Nyctalus_leisleri | -0.066828 | -0.620133 | 0.508768 |
| Nyctalus_lasiopterus - Rhynchonycteris_naso | -1.9101461 | -2.699912 | -1.113829 |
| Nyctalus_lasiopterus - Thyroptera_tricolor | -1.120585 | -2.043498 | -0.2335453 |
| Nyctalus_leisleri - Rhynchonycteris_naso | -1.8438125 | -2.536709 | -1.16040616 |
| Nyctalus_leisleri - Thyroptera_tricolor | -1.054798 | -1.85393 | -0.226147 |
| Rhynchonycteris_naso - Thyroptera_tricolor | 0.788069 | -0.189839 | 1.772014 |

**Table S7.** Results for 95% pairwise highest posterior density (HPD) intervals between bats classified into long distance, regional, or non-migrants among three different genetic measures. The genetic measures are population differentiation, gene diversity, and allelic richness.

| **Population Genetic Measure** | **Contrast** | **Estimate** | **Lower.HPD** | **Upper.HPD** |
| --- | --- | --- | --- | --- |
| Population Differentiation | long_distance_migrant - non_migratory | 0.001 | -0.074 | 0.075 |
|  | long_distance_migrant - regional_migrant | -0.002 | -0.069 | 0.064 |
|  | non_migratory - regional_migrant | -0.003 | -0.049 | 0.046 |
| Gene Diversity | long_distance_migrant - non_migratory | 0.054 | -0.154 | 0.266 |
|  | long_distance_migrant - regional_migrant | 0.111 | -0.069 | 0.296 |
|  | non_migratory - regional_migrant | 0.057 | -0.098 | 0.215 |
| Allelic Richness | long_distance_migrant - non_migratory | 0.441 | -1.453 | 2.411 |
|  | long_distance_migrant - regional_migrant | 0.821 | -0.879 | 2.480 |
|  | non_migratory - regional_migrant | 0.372 | -1.122 | 1.796 |

**Table S8.** Results for 95% pairwise highest posterior density (HPD) intervals between bats classified into harem or non-harem species among three different genetic measures. The genetic measures are population differentiation, gene diversity, and allelic richness.

| **Population Genetic Measure** | **Contrast** | **Estimate** | **Lower.HPD** | **Upper.HPD** |
| --- | --- | --- | --- | --- |
| Genetic Differentiation | harem - (non-harem) | -0.01235035 | -0.0644466 | 0.0409534 |
| Gene Diversity | harem - (non-harem) | 0.0204414 | -0.162776 | 0.205463 |
| Allelic Richness | harem - (non-harem) | 0.0783008 | -1.55628 | 1.71139 |

**Table S9**. Estimated differences (Est.), error (Est. Error), lower and upper 95% credible intervals (Lower CI, Upper CI, respectively), evidence ratios (Evid. Ratio), and posterior probabilities (Post. Prob.) for specific contrasts from different models. The models are categorized by population genetic measure and behavioural trait. For example, the model of genetic differentiation and mating strategy is abbreviated as “Genetic Differentiation, Mating”. Contrasts specify the specific groups and direction of difference explored within each model. For example, the results for the test exploring whether non-harem mating bats have higher genetic differentiation than those with harems is specified by the contrast “Non-Harem > Harem”.

| **Model** | **Contrast** | **Est.** | **Est. Error** | **Lower CI** | **Upper CI** | **Evid. Ratio** | **Post. Prob.** |
| --- | --- | --- | --- | --- | --- | --- | --- |
| Genetic Differentiation, Mating | Non-Harem > Harem | 0.01 | 0.02 | -0.03 | 0.05 | 2.61 | 0.72 |
| Gene Diversity, Migration | Long Distance Migrant > Regional Migrant | 0.1 | 0.09 | -0.04 | 0.24 | 7.53 | 0.88 |
| Gene Diversity, Migration | Long Distance Migrant > Non-Migrant | 0.04 | 0.10 | -0.13 | 0.2 | 1.86 | 0.65 |
| Allelic Richness, Migration | Long Distance Migrant > Regional Migrant | 0.58 | 0.74 | -0.62 | 1.79 | 3.84 | 0.79 |
| Allelic Richness, Migration | Long Distance Migrant > Non-Migrant | 0.09 | 0.86 | -1.3 | 1.5 | 1.19 | 0.54 |
